# Essential role of translation factor eIF5a in cytokine production and cell cycle regulation in primary CD8 T lymphocytes

**DOI:** 10.1101/2021.06.25.449879

**Authors:** Thomas CJ Tan, Van Kelly, Xiaoyan Zhou, Tony Ly, Rose Zamoyska

## Abstract

Translational control adjusts protein production rapidly and facilitates local cellular responses to environmental conditions. Translation can be regulated through sequestration of mRNAs by regulatory proteins or RNAs, but also by the availability of ribosomes and translation factors which enable initiation and elongation of nascent polypeptides. Traditionally initiation of mRNA translation has been considered to be a major translational control point, however, control of peptide elongation can also play a role. Here we show that post-translational modification of the elongation factor, eIF5a, controls translation of subsets of proteins in naïve T-cells upon activation. Sequencing of nascent polypeptides indicated that functional eIF5a was required for the production of proteins which regulate T-cell proliferation and effector function. Control of translation in multiple immune cell lineages is required to co-ordinate immune responses and these data illustrate that translational elongation can contribute to post-transcriptional regulons important for the control of inflammation.

## INTRODUCTION

Regulation of protein expression at the level of ribosome production and translation can rapidly rewire and co-ordinate responses to changing environmental cues. Poor correlations between transcriptomes and proteomes have been documented in numerous cell types, including stem cells (Munoz et al., 2011) and T lymphocytes (Hukelmann et al., 2015) indicating that a substantial proportion of their protein production is regulated translationally. Both these lineages can undergo rapid expansions and changes in cell identity in response to environmental triggers (Gabut et al., 2020). Regulation of ribosome biogenesis and global mRNA translation (Gabut et al., 2020) contribute to this plasticity together with more targeted post-transcriptional regulons, important for directing the production of proteins with related functions or involved in common biological pathways (Anderson, 2010). Regulation of such subsets of mRNAs frequently involves interactions with RNA-binding proteins or non-coding RNAs (eg miRNA and lncRNA) (Ciafrè and Galardi, 2013; Zhang et al., 2019b). In addition alterations in ribosome constituents or the proteins involved with the translation of mRNAs may play a role. For example, we showed that for naïve CD8 T cells the degree of ribosome biogenesis was dramatically affected by the extent of activation through the T cell receptor (TCR), which impacted global translation and also selectively affected production of some groups of proteins over others (Tan et al., 2017). How this plasticity of translational control is orchestrated around the ribosome is not well understood, but a number of proteins are directly involved with the ribosome during translation (Dever et al., 2018; Sonenberg and Dever, 2003) and these may themselves be regulated both translationally and post-translationally.

One such protein is the translation factor eIF5a, one of the top 20 most abundant proteins in CD8 T cells (Hukelmann et al., 2015). eIF5a was originally identified as a translation initiation factor, however recent work in yeast has indicated a role in translation elongation and termination (Schuller et al., 2017). Depletion of eIF5a in *S. cerevisiae* reduced protein synthesis by approximately 30%, raising the possibility that eIF5a is required for translation of subsets of proteins, rather than being a ubiquitous translation factor (Kang and Hershey, 1994). *In vivo* reporter and *in vitro* translation assays revealed that eIF5a is required for efficient peptide bond formation between consecutive proline residues, a function described for its bacterial homolog, EF-P (Gutierrez et al., 2013). Two independent yeast ribosome profiling studies found the requirement for eIF5a is not limited to elongation between poly-proline sequences, but that it also influences translation of glycine and charged amino acids, suggesting a wider involvement of eIF5a in regulating translation of mRNAs with common motifs, (Pelechano and Alepuz, 2017; Schuller et al., 2017) and suggesting it might be an important player in influencing post-transcriptional regulons.

eIF5a isoforms in eukaryotes and archaea are the only proteins with post-translational attachment of a hypusine residue on Lys^50-51^, a modification that is essential for eIF5a function (Park and Wolff, 2018). Hypusination involves enzymatic transfer of the 4-aminobutyl moiety of the polyamine spermidine to lysine, catalysed by deoxyhypusine synthase (DHPS) to produce the deoxyhypusinated intermediate form, and subsequent hydroxylation by deoxyhypusine hydroxylase (DOHH) to form the mature hypusinated protein. The site of hypusination is in contact with the acceptor stem of the P site tRNA on 80S ribosome (Schmidt et al., 2016), and it is postulated that hypusination is required for eIF5a activity as non-modified eIF5a is inefficient in facilitating peptide synthesis (Gutierrez et al., 2013) and inhibition of the hypusination pathway impairs proliferation in mammalian cell lines (Colvin et al., 2013; Hanauske-Abel et al., 1994). Further evidence for the importance of eIF5a hypusination comes from studies of spermidine, a natural polyamine which is the donor moiety for eIF5a hypusination and whose abundance is reduced in ageing cells. Specifically, in T cells spermidine dietary supplementation restored both the abundance of eIF5a hypusination and age-related impairments in T cell function (Puleston et al., 2014). Spermidine administration also extended the life span of yeast, flies, worms, and cultured human PBMCs (Eisenberg et al., 2009). Therefore strong evidence links the abundance of eIF5a hypusination and its function in age-related immunosenescence, although it is currently unclear precisely how these processes are associated.

The aim of this study was to determine firstly how the activity of eIF5a itself was regulated in T lymphocytes, and secondly, whether eIF5a has a selective role in regulating translation of specific subsets of mRNA during T cell activation. We used CRISPR knockout of eIF5a itself and of the key enzymes, DHPS and DOHH, involved in adding the hypusine modification to eIF5a and compared the effect of disrupting the latter to inhibition of hypusination by the deoxyhypusine synthase inhibitor, GC7. We show that although naïve T cells express abundant eIF5a, the degree of hypusine modification of the eIF5a protein is relatively low suggesting limited functionality. Upon activation of T cells, eIF5a protein production is upregulated and DHPS and DOHH enzymes are expressed rapidly so that the hypusinated form of eIF5a becomes more abundant. Total loss of eIF5a surprisingly increased production of some ribosomal proteins and eIF6, a possible translational inhibitor, while decreasing some translation initiation factors. Together these results suggested that alterations in ribosome composition occur in the absence of eIF5a. Mass spectrometry analysis of nascent peptide production in eIF5a knockout T cells confirmed unequal impacts on synthesis of specific sets of proteins in the absence of eIF5a, particularly those involved in cell cycle regulation and cytokine production. We substantiated that loss of functional eIF5a resulted in a specific loss of proteins containing polyproline motifs in primary T cells, as described previously for yeast cells (Gutierrez et al., 2013). However treatment with GC7 did not faithfully reproduce the phenotypes of knockout cells indicating a significant off-target impact of this drug. Overall our data are consistent with a role for eIF5a in co-ordinating post-transcriptional regulons in T lymphocytes and reveal how these may influence primary T cell function.

## RESULTS

### T cell activation enhances production and functional maturation of eIF5a

A unique feature of T cells is that they exist for long periods as naïve cells which are functionally immature, non-dividing and metabolically inert, with very low levels of macromolecular synthesis (Wolf et al., 2020). Upon stimulation through their antigen specific TCRs they rapidly turn on gene transcription and protein synthesis. In the initial 24h following stimulation CD8 T cells increase cytoplasmic mass by approximately 4-fold, indicating a rapid initiation of macromolecular synthesis, before commencing a burst of proliferation with a division time as short as 2-4h (Lawrence and Braciale, 2004; Yoon et al., 2007).

We examined the functional maturation of eIF5a in OT-1 CD8 T cells which express a single specificity transgenic TCR that recognises a peptide from chicken ovalbumin (SIINFEKL) presented by the MHC allele, H-2D^b^. Western blot of whole cell lysates from naïve unstimulated OT-1 T cells showed abundant expression of eIF5a, but low levels of staining with a hypusine specific Ab (Fig 1A and B). Upon stimulation of the T cells with peptide, the abundance of hypusinated eIF5a increased by ~3-fold, concurrent with a similar increase in expression of DHPS, the enzyme that catalyses transfer of the 4-aminobutyl moiety of spermidine to eIF5a. The abundance of both hypusinated eIF5a and DHPS peaked between 24-48h after stimulation and increased by approximately two-fold above that of a control protein, ZAP-70. Single cell measurement of hypusinated and total eIF5a using flow cytometry (Fig 1C) confirmed these data, showing around a 7-fold increase in hypusination and 4-fold increase in total eIF5a after 24h of activation, in comparison to naïve ex vivo T cells (0h). Data from Western blot and flow cytometry measurements confirmed that the proportion of hypusinated eIF5a to total eIF5a was consistently increased by 1.5- to 2-fold within 48h of activation.

**Fig 1.**
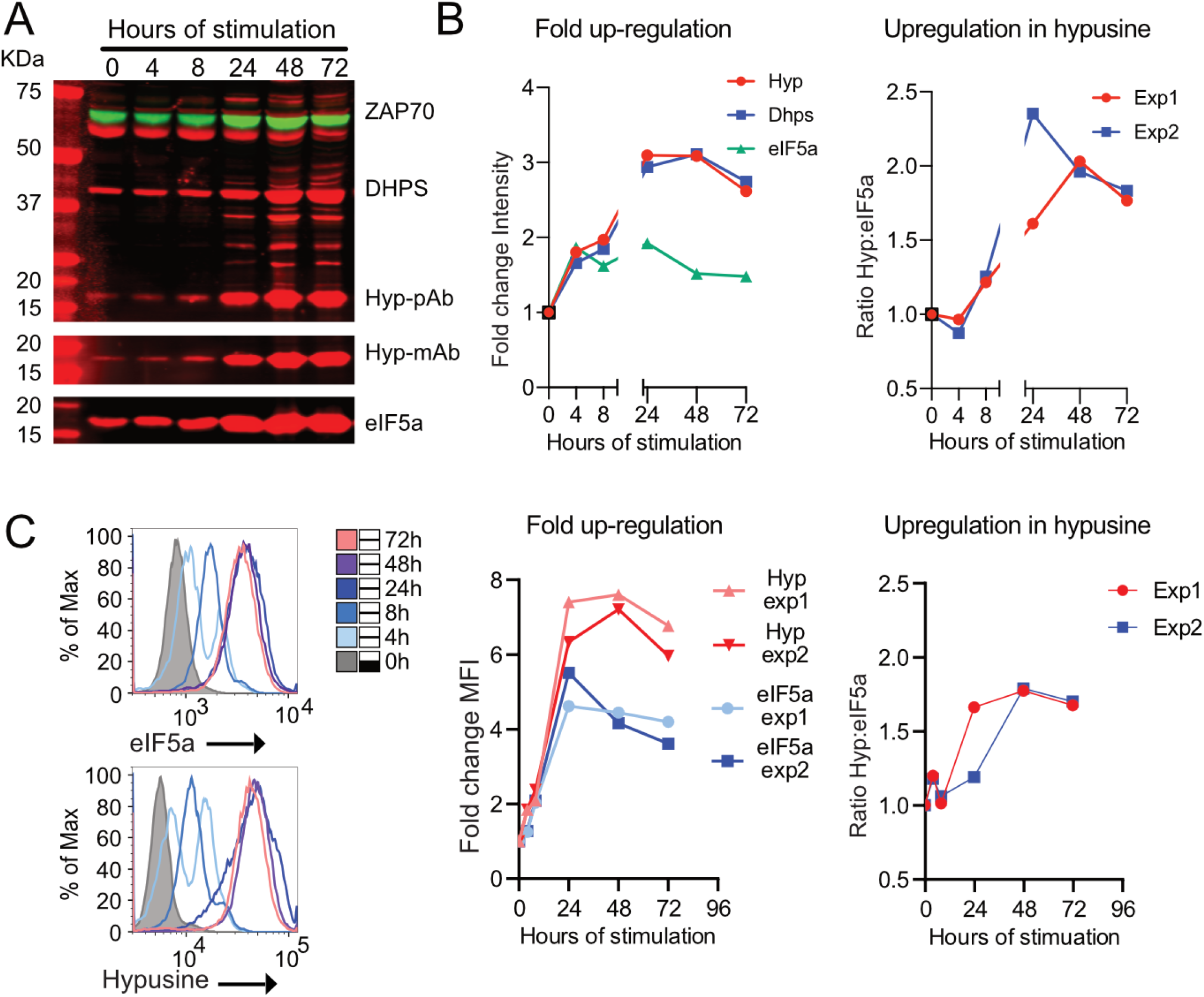
Hypusination of eIF5a is induced by T cell activation. (A) Naïve OT-1 lymph node CD8 T cells were stimulated with agonist peptide SIINFEKL for the times indicated and western blot of total cell lysates were probed with Abs for DHPS, hypusine, eIF5a and ZAP70 as control, equal cell numbers were loaded in each lane. (B) Fold up-regulation and Hyp:eIF5a ratio were calculated relative to the 0h time point, after normalisation using the intensity of ZAP70 at the same time point. Representation of three independent experiments is shown. (C) Formaldehyde-fixed OT-1 cells following the same stimulation time course was permeablised, stained with Abs against eIF5a and hypusine and the correspondent secondary antibody, and measured using flow cytometry. Relative mean fluorescent intensities (MFI) from two independent experiments are shown and were used to calculate the Hyp:eIF5a ratio on the right.

### Abrogation of eIF5a or its maturation pathway differentially disrupts key cellular functions

Hypusination of eIF5a allows efficient facilitation of translation elongation *in vitro* (Gutierrez et al., 2013) and may be responsible for its cytoplasmic localisation (Lee et al., 2009). However it is unclear whether or not the non-hypusinated precursor and deoxyhypusinated intermediate forms of eIF5a have any activity in primary cells. This question is important as it is uncertain whether targeting DHPS and DOHH as means to inhibit eIF5a are optimal strategies, given deletion of budding yeast DOHH had only a mild phenotype (Park et al., 2006) whereas embryonic targeting of DOHH in mouse was lethal (Sievert et al., 2014). Moreover most studies of eIF5a function utilise the drug GC7 to inhibit hypusination and, therefore, function of eIF5a both *in vivo* and *in vitro*. GC7 is a spermidine derivative which competes with spermidine for binding to DHPS, thereby preventing the conversion of non-hypusinated eIF5a precursor to deoxyhypusinated intermediate, but it is undocumented whether GC7 has effects beyond inhibiting eIF5a.

In order to address these questions, naïve OT-1 cells were activated for 2 days, and then either treated with GC7 for 2 days, or transfected with CRISPR-Cas9 guides to disrupt *eIF5a* or *Dhps* or *Dohh* genes. KO cells were recovered after 3 days (*eIF5a* KO) or 4 days (*Dhps* and *Dohh* KO), with the duration of the culture adjusted for the different KOs to maximise viable cell recovery. CRIPSR-Cas9 transfection controls were targeted for the unrelated surface molecule, Thy 1. Given targeting efficiency was maximally 80%, both gene KO and untargeted cells from the same culture could be compared directly in the analysis by using intracellular staining for eIF5a or hypusine to identify positive cells. Fig 2A shows a representative experiment (n≥7) with a mean percentage KO of *Eif5a*, *Dhps*, and *Dohh* of 38.4%, 71.6% and 78.9%, respectively. eIF5a depletion is known to affect cellular proliferation (Kang and Hershey, 1994; Mathews and Hershey, 2015), and this likely accounts for the apparently reduced efficiency of targeting of eIF5a itself, as KO cells will be at a proliferative disadvantage relative to untargeted cells in the same culture. There was a decrease also in the number of cells recovered from cultures in which hypusination was impaired, including GC7-treated and *Dhps* and *Dohh* KOs relative to control populations, but this was not as profound as for the *Eif5a* KO.

**Fig 2.**
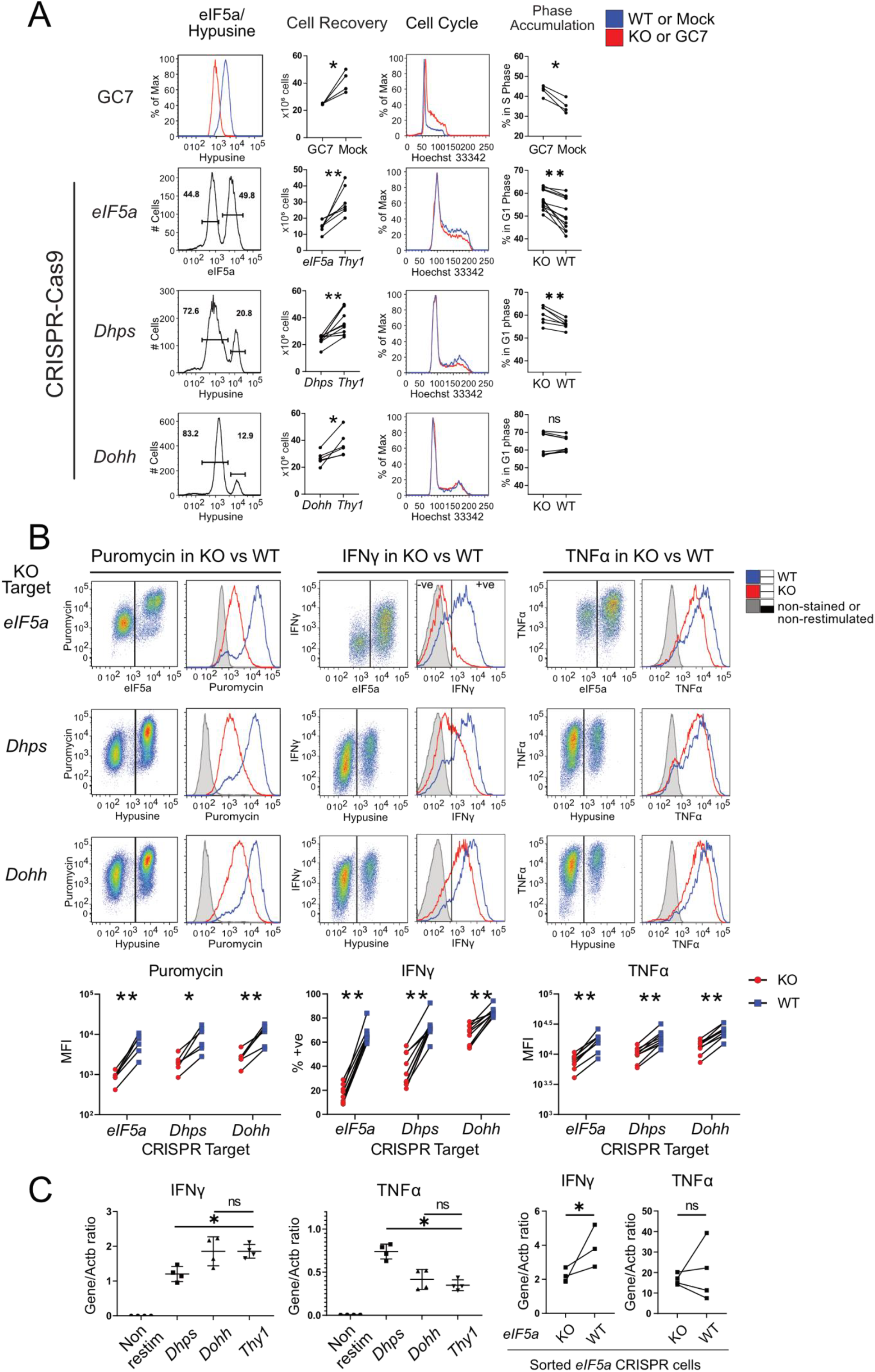
eIF5a selectively influences protein synthesis and cytokine production in CD8 T cells. (A) OT-1 T cells were stimulated for 48h before CRIPR KO of eIF5a, DHPS or DOHH or treatment with 20μM GC7. Expression of eIF5a or hypusine was monitored after further 2d (GC7), 3d (eIF5a) and 4d (DHPS and DOHH) of culture in IL-2. Cell cycle profiles were obtained by Hoechst 33342 staining. Cell recovery and percentage of cells in the cell cycle phase (S phase in GC7-treated cells and G1 phase in others) were plotted, pairing positive and negative cells from individual cultures (n ≥ 4, *. P < 0.05, **. P < 0.01, paired T test). KO of cell surface marker Thy1 was included as an irrelevant CRISPR control. (B) Cells were transfected and cultured as above and re-stimulated with peptide for 4h or not (controls). Puromycin incorporation, IFNγ and TNFα, were measured using flow cytometry. Lower panel: Mean fluorescent intensity (MFI) of puromycin and TNFα, and percentage of IFNγ-positive cells from >6 experiments were plotted (* p < 0.05, ** p< 0.01, paired T-test). (C) RT-qPCR detection of mRNA for IFNγ and TNFα in bulk transfected cells as indicated (n= 4, * p< 0.05, unpaired T test) or fixed sorted cells for eIF5a KO (n ≥ 3, * p < 0.05, paired T test).

To further characterise any proliferation defect, cell cycle analysis of was performed by staining with Hoechst 33342. Surprisingly GC7 inhibition of eIF5a hypusination resulted in cell cycle profiles that were distinct from those observed following knockout of *Eif5a*, *Dhps*, or *Dohh*. Fig 2A shows a pronounced cellular accumulation in S phase following GC7 treatment which was less apparent in the KOs and suggests possible GC7 off-target effects beyond its impact on eIF5a hypusination. Knockout of *Dohh* did not result in noticeable changes in the cell cycle profile despite negatively impacting hypusination and cell numbers and suggested that the deoxyhypusinated intermediate form of eIF5a is partially active, possibly slowing progression through rather than arresting the cell cycle.

Cells lacking eIF5a, DHPS or DOHH were further analysed for their protein synthesis rates and cytokine production. After CRISPR-Cas9 transfection and maintenance in IL-2 cytokine, the cells were re-stimulated with peptide for 4h, in order to reactivate effector cytokine transcription and translation, and labelled with puromycin during the last 20 minutes. Incorporation of puromycin into nascent proteins can be analysed by flow cytometry and used to assess the rate of global protein synthesis. In addition, the production of specific CD8 T cell effector proteins, IFNγ and TNFα were measured by intracellular staining (Fig 2B). *Eif5a* KO cells reduced puromycin incorporation by ~84% while there was slightly less reduction in protein synthesis compared to untargeted cells in the *Dhps* and *Dohh* KO cells (n=6). Cytokine production showed dramatic decrease in IFNγ production in *Eif5a* KO cells, with the majority of the population falling in the negative gate (Fig 2B). In contrast the reduction in IFNγ production was less pronounced in *Dhps* KO cells (~60% negative cells), while *Dohh* KO cells showed only a slight reduction in IFNγ (~25% negative cells). Interestingly the loss of IFNγ production was selective rather than reflecting a general shut down in cytokine translation, as all the targeted cells showed a much smaller, albeit significant, reduction in TNFα production.

To determine at what stage of production these cytokines were regulated the abundance of IFNγ and TNFα mRNA was assessed. RT-qPCR was performed on cell lysates from unseparated *Thy1*, *Dhps* and *Dohh* targeted populations given the frequency of KO cells was routinely ≥80%. As the *Eif5a* targeted populations were closer to 50% of the total, the population was fixed for intracellular eIF5a staining and sorted into targeted (KO) and control (WT) subsets from the same culture (Fig 2C). In contrast to the non-restimulated control cells, all cells restimulated with peptide significantly upregulated IFNγ and TNFα mRNA as expected. Compared to their untargeted or *Thy-1* targeted control cells, IFNγ mRNA was decreased significantly in *eIF5a* KO and *Dhps* KO cells, but not in the *Dohh* targeted population. No significant reduction in TNFα mRNA abundance was observed in any population, curiously however, in the absence of DHPS TNFα mRNA was increased significantly. Taken together, these results indicate that lack of fully matured eIF5a does not impair TNFα mRNA transcription and has a very mild effect on its translation. By contrast, mature eIF5a is required both for transcription and translation of IFNγ. However, it is unclear whether this effect is direct, or else an indirect consequence of the loss of an eIF5a-dependent transcription factor.

### Systematic analysis of eIF5a-regulated translation in CD8 T cells

To ask whether specific groups of proteins were regulated by functional eIF5a, global nascent peptide profiling by incorporation of 4-Azido-L-homoalanine (AHA) into nascent peptides in place of methionine was performed on *eIF5a* KO and GC7-treated T cells. Cultures were pulsed with AHA for 2h and *eIF5a* KO and WT populations were separated by cell sorting for eIF5a following fixation and staining, ensuring that untargeted WT cells were subjected to identical culture and labelling conditions as the KO cells. Following lysis of the sorted cells, AHA-labelled peptides were isolated using copper-mediated Click chemistry with Alkyne agarose beads and purified nascent proteomes were identified and quantified using LC-MS.

Single shot label-free proteomics of 12 fractions of pooled GC7- and Mock-treated cells resulted in detection of 7529 proteins which served as the library for match-between-runs in MaxQuant. Using this library, approximately 5500 proteins were identified as significantly higher than the background (at least 5 times more abundant in the AHA-labelled samples compared to non-labelled controls), compared to 4081 proteins outwith the library. Four biological replicate samples were used for each group (Fig 3A, left). iBAQ values were used to measure absolute quantities of protein and to compare between groups (Schwanhäusser et al., 2011). The iBAQ values were normalised by the ratio of total iBAQ in each sample to the mean total iBAQ within the same group to minimise variations caused by loading and acquisition, and then normalised again by the input cell number to allow comparison between groups (Fig 3A, right; Fig S1A-B; Table S1). Amongst the four sample groups (Mock, GC7, eIF5a KO and eIF5a WT), iBAQ values in GC7-treated cells appeared to be globally reduced by ~3-fold compared to Mock-treated cells, this global decrease in AHA incorporation was consistent with previous trial proteomic experiments using various ratios between GC7- and Mock-treated cells (Fig S1C), and therefore is unlikely to be caused by the input normalisation. 3D MDS plot of all nascent proteins detected between *eIF5a* KO and WT cells and between GC7-treated and mock-treated control cells (Fig 3B) showed distinctive signatures separating the 4 groups with the largest source of variations (Dimension 1) separating KO/WT from GC7/Mock, most likely due to formaldehyde fixation and the subsequent reverse crosslinking in the KO/WT samples. The second largest source of variation (Dimension 2) separated KO from WT and GC7 from Mock and is most likely caused by the manipulation of eIF5a activity. Differential expression analyses were conducted between GC7- and mock-treated cells and between eIF5a KO and WT cells. GC7 had a systematic effect on protein production, with 3044 proteins down-regulated and only 31 proteins up-regulated (Fig S2A); while eIF5a KO had a more limited effect on protein production, with 369 down-regulated and 149 up-regulated proteins (Fig S2B).

**Fig 3.**
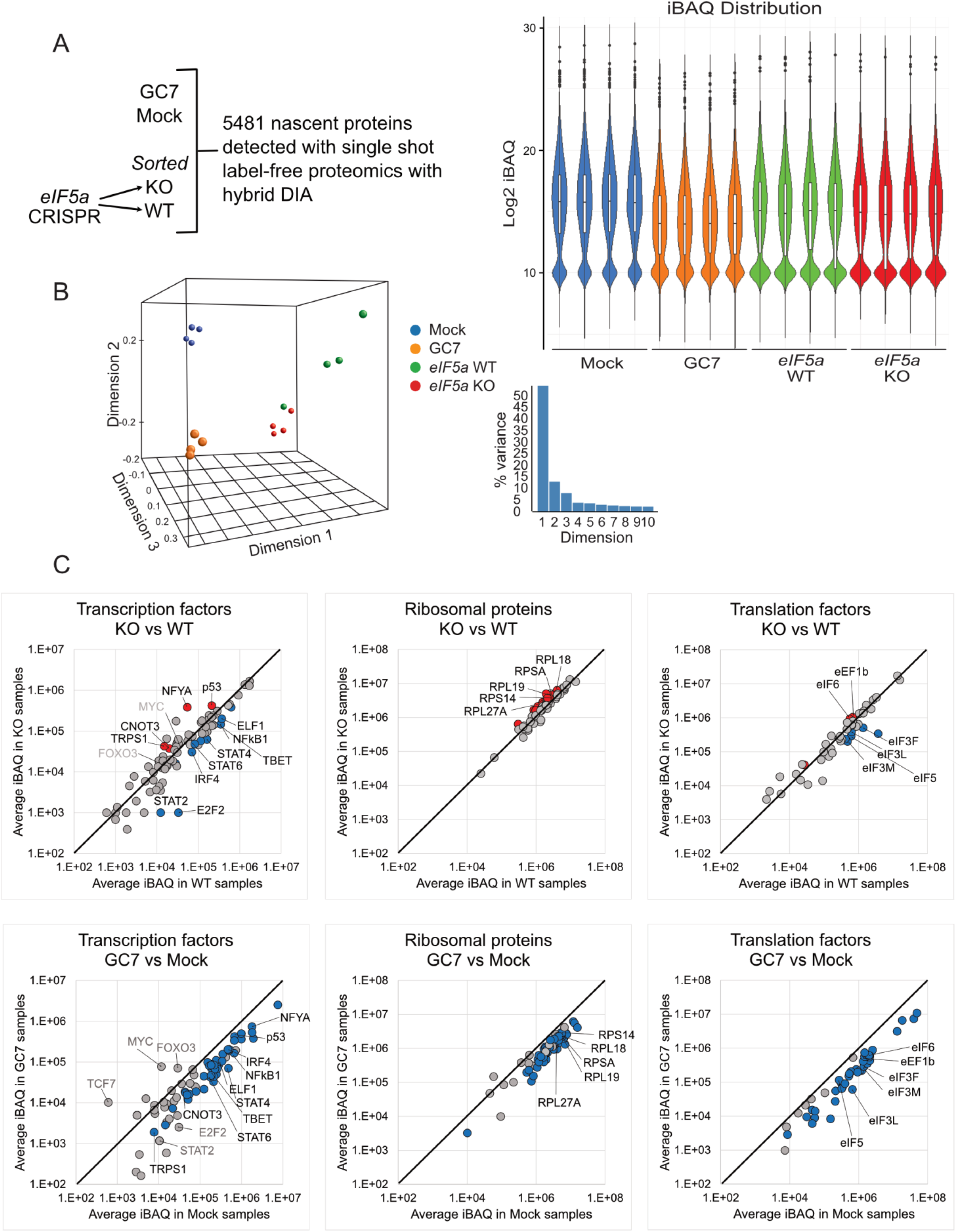
Systematic analysis of GC7- or eIF5a KO-regulated nascent proteomes. (A) OT-1 T cells were stimulated with peptide for 48h before CRIPR KO of eIF5a or incubation with GC7 for a further 48h, as before. Cells were labelled with AHA for the final 2h of culture. eIF5a CRISPR KO cells and WT controls were obtained by FACS sorting. AHA-labelled proteins were isolated by Click-chemistry with alkyne agarose beads and digested with trypsin. In total 5481 proteins (left) were significantly more abundant in AHA-labelled samples compared to unlabelled controls. (Right) iBAQ values from each group were normalised by the mean summary of each sample within the group, and then by the input cell number. Log2-transformed values in each sample after normalisation were plotted as violin plots. (B) 3D-MDS plot was used to visualise the difference between samples (left). The percentage of variance represented by each dimension is plotted on the right. (C) Average normalised protein iBAQ values for all detected transcription factors, ribosomal proteins, and translation factors were compared between KO and WT cells or between GC7- and mock-treated cells. Significantly (P value or FDR < 0.05, Absolute FC > 1.5) changed proteins of interest are annotated in black font while proteins not deemed significant by the statistical test, but may still be biologically relevant are annotated in grey font.

Clustering of GO terms on molecular functions was performed (WebGestalt (Liao et al., 2019)) on the down- and up-regulated proteins in each comparison, compared to all MS-detected newly synthesised proteins as the background (Fig S2). Noticeably transcription factors, ribosomal proteins and translation factors were affected by both GC7 and *Eif5a* KO but in different ways (Fig 3C). Amongst all detected transcription factors from the mass spec data, notably production of E2F2 was profoundly impacted in *Eif5a* KO cells, which correlated with their impaired proliferation. Several key regulators of environmental response and cytokine production, including IRF4, RELA, NFκB1, STAT2, STAT4, STAT6, and TBET (Tbx21) were also downregulated in their production, with STAT2 being the most impacted. Some transcription factors were upregulated, such as the tumour suppressor p53, CNOT3 and TRPS1 which may control cell cycle (Shirai et al., 2019; Wu et al., 2014) and NFYA, whose function in T cells is undocumented.

Given the profound effect of *Eif5a* KO on puromycin incorporation, we investigated components of the cellular translation machinery in the nascent proteome. Amongst the 73 detected ribosomal proteins, 11 were up-regulated in *Eif5a* KO cells (FC>1.5, P<0.05, paired T test, n=4) while none was down-regulated. Up-regulation of specific ribosomal proteins might suggest that ribosomes with alternative composition were being produced in cells in the absence of eIF5a and these ribosomes may preferentially translate a different subset of mRNAs compared to those in WT cells (Segev and Gerst, 2018; Shi et al., 2017). Additionally, down-regulation of translation factors eIF1a, eIF3 components and eIF5 suggest the formation and function of the 43S pre-initiation complex may be reduced (Jackson et al., 2010), while an increase in eIF6 would prevent the 60S ribosome subunit from joining with the 40S subunit (Ceci et al., 2003). Together these data illustrate a general reduction in protein production, and an altered translation profile in *Eif5a* KO cells. All the aforementioned proteins were down-regulated upon GC7 treatment, illustrating the systematically inhibited nature of protein synthesis in these samples. Polyamines are known to modulate both the efficiency and fidelity of protein synthesis, by direct interaction with the ribosome (Dever and Ivanov, 2018). It is possible that the eIF5a-independent effects of GC7 may result from its competitive binding to ribosomes and thus displacing other polyamines present in the cell.

To further separate direct effects of eIF5a on translation from indirect effects following changes in transcription factor production, next generation transcriptomic sequencing of GC7- and Mock-treated samples was performed (Table S2). Although GC7-treatment has off target effects independent of eIF5a, the analysis was performed on these populations as the quality of RNA extracted from these fresh cells was suitable for transcriptomic analyses, whereas RNA extracted from eIF5a KO cells, which needed to be formaldehyde-fixed for sorting, was not.

Translationally down-regulated genes in the GC7 treated samples were defined as the 1635 genes with no significant reduction in RNA abundance (FDR>0.1, edgeR, n=4) but decreased in their nascent proteins (Fig 4A left Venn diagram and Table S3, FDR<0.05, FC<−1.5, unpaired T test with Benjamini-Hochberg correction, n=4). These genes were overlaid with those reduced in their nascent proteins in eIF5a KO compared to WT cells (Fig 4A, right Venn diagram and Table S3, P<0.05, FC<−1.5, paired T test, n=4) which gave 119 genes whose translation, rather than transcription, we could confidently assign as being dependent on eIF5a. Proteins translationally down-regulated only by GC7 treatment were enriched in GO terms “structural constituent of ribosome”, “mRNA binding”, “rRNA binding”, and “translation factor activity” (FDR < 0.01); while the 119 proteins translationally reduced by both GC7 treatment and eIF5a KO did not significantly cluster into any GO terms, although some key genes regulating cellular functions such as mTORC1 modulators SLC3A2 and SLC7A5, cell cycle regulators Cyclin B1, CDK1 and RFC3/5, MHC Class I genes H2-D1, H2-K1 and H2-Q4, transcription factors FOXP1, RELA, IRF4 and TBET, and translation factors eIF3F/L/M were in this Venn group (Table S3).

**Fig 4.**
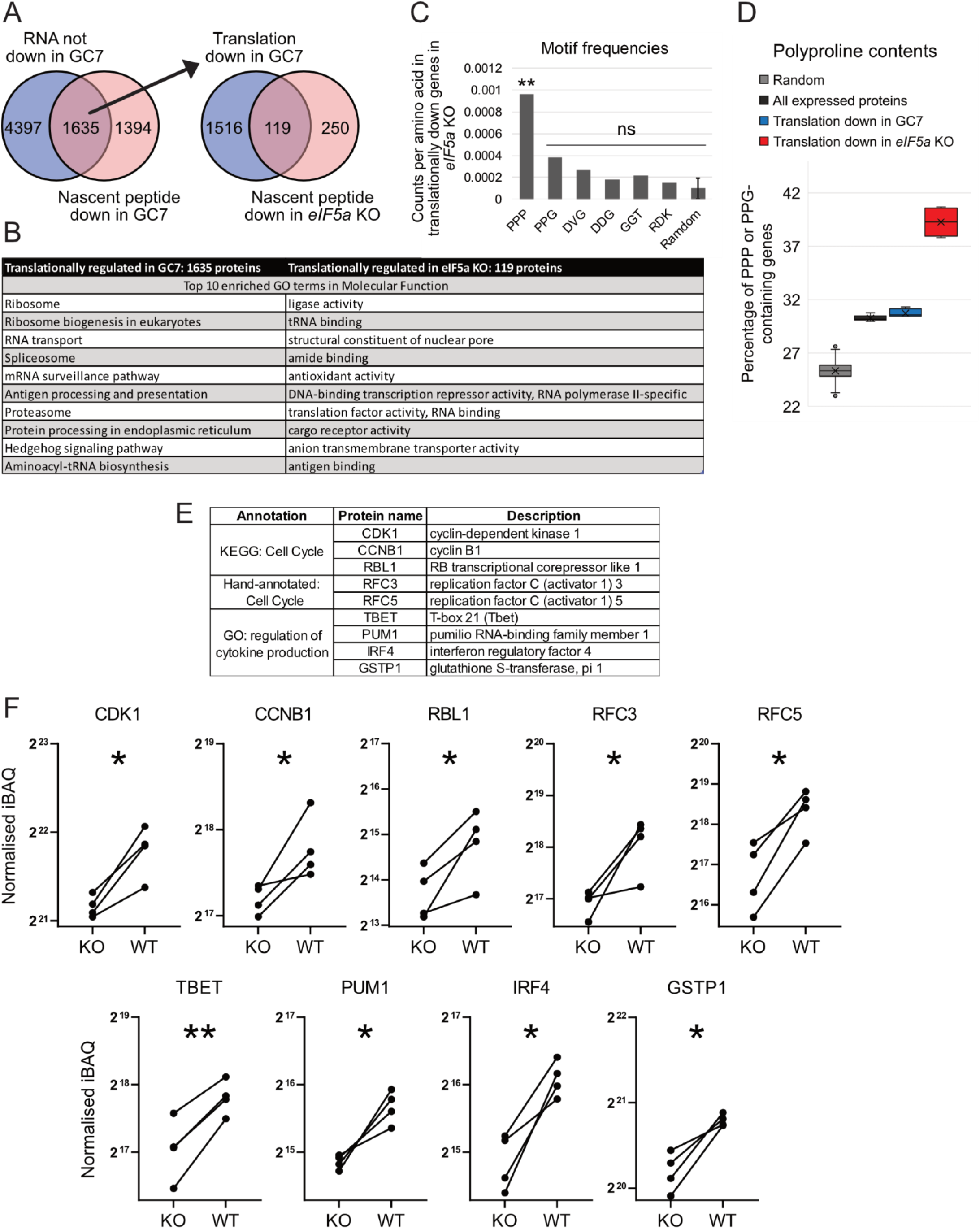
Identification of translationally regulated genes affected by GC7 treatment and eIF5a KO. (A) Venn diagram overlays identify 1635 of proteins unchanged in their mRNA (FDR>0.05, edgeR, n = 4) following GC7 treatment but downregulated in their nascent peptides (FDR<0.05, FC<−1.5, unpaired T test with Benjamini-Hochberg correction, n=4). These proteins were further overlaid with those reduced in their nascent proteins in eIF5a KO cells (P<0.05, FC<−1.5, paired T test, n = 4) as an extra criterion of selection for proteins translationally regulated by eIF5a (119 proteins). Lists of genes in these Venn groups are in Table S3. (B) Top ten enriched GO terms in molecular function were listed for proteins translationally affected by GC7-treatment or by eIF5a KO. (C) Frequencies of specific 3mer sequence motifs in the eIF5a-regulated gene set, normalised by the total length of these proteins. Mean and standard deviation of normalised counts of 50 randomly generated 3mer sequences is plotted at the right (**. P<0.01, one-sided Fisher’s exact test). (D) Plots of the percentage of PPP or PPG-containing genes in 2900 randomly selected genes in the transcriptome (repeated 1000 times), all expressed proteins, translationally down-regulated genes in GC7-treated or eIF5a KO cells in 4 biological samples. (E) Selected eIF5a-regulated proteins involved in regulation of cell cycle progression and cytokine production from the datasets are highlighted and (F) their normalised iBAQ values for nascent proteins are plotted for eIF5a KO and WT cells (*. P<0.05, FC<−1.5, paired T test, n = 4).

Given the well studied role of eIF5a in translation through polyproline and charged amino acid regions of proteins, the frequencies of previously characterised eIF5a-controlled 3-mer motifs (PPP, PPG, DVG, DDG, GGT, and RDK (Schuller et al., 2017)) were counted in the coding sequence (CDS) of the eIF5a-regulated gene set (Fig 4C). Compared to the frequencies of 50 randomly selected 3-mer sequences, PPP is enriched (P<0.01, one-sided Fisher’s exact test). PPG and DVG were possibly enriched but rejected by the statistic test (P>0.99). To assess the polyproline content of these genes, they were overlaid with a published list of genes in the mouse genome with PPP or PPG motifs in their coding region (Mandal et al., 2014). We plotted the percentage of genes containing polyproline in 2900 randomly selected genes in the transcriptome (repeated 1000 times), all expressed proteins detected by LC-MS, and translationally down-regulated genes in GC7-treated or *eIF5a* KO cells in 4 biological samples shown in Fig 4D. In GC7-treated cells, the percentage of polyproline genes was similar between all expressed and translationally down-regulated proteins, suggesting no enrichment for polyproline while the translationally down-regulated proteins in *eIF5a* KO cells had a noticeable increase in polyproline content (Fig 4D), consistent with a more targeted inhibition of eIF5a in the KO versus GC7-treated samples.

We then focused our analyses and validation on proteins involved in regulating cytokine production and cell cycle progression whose translation we had identified as regulated by the availability of mature eIF5a (listed in Fig 4E-F), since defects of these cellular functions were observed in *Eif5a* KO cells. Moreover, these cellular functions are the major age-related impairments described in CD8 T cells (Po et al., 2002), and so might illustrate a possible role eIF5a plays in immunosenescence. Of these genes, we were able to obtain suitable antibodies against IRF4, TBET and CDK1 for flow cytometry which when stained together with antibody against eIF5a or hypusine allowed visualisation of these proteins in the KO and WT populations within the same culture (Fig 5A). Both IRF4 and TBET are transcription factors regulating production of IFNγ and TNFα in T cells (Raczkowski et al., 2013; Szabo et al., 2002), and CDK1 interacts with different cyclins to facilitate cell cycle progression (Santamaría et al., 2007). In agreement with the systematic datasets, the abundance of IRF4 dropped in *Eif5a* KO, *Dhps* KO and to a lesser extent in *Dohh* KO cells, with no reduction in its RNA levels in these cells (Fig 5B). Similarly, TBET and CDK1 proteins decreased in *Eif5a* KO and *Dhps* KO, but not in *Dohh* KO cells, in correlation with their cell cycle profiles in Fig 2A, and again their RNA abundances were unchanged.

**Fig 5.**
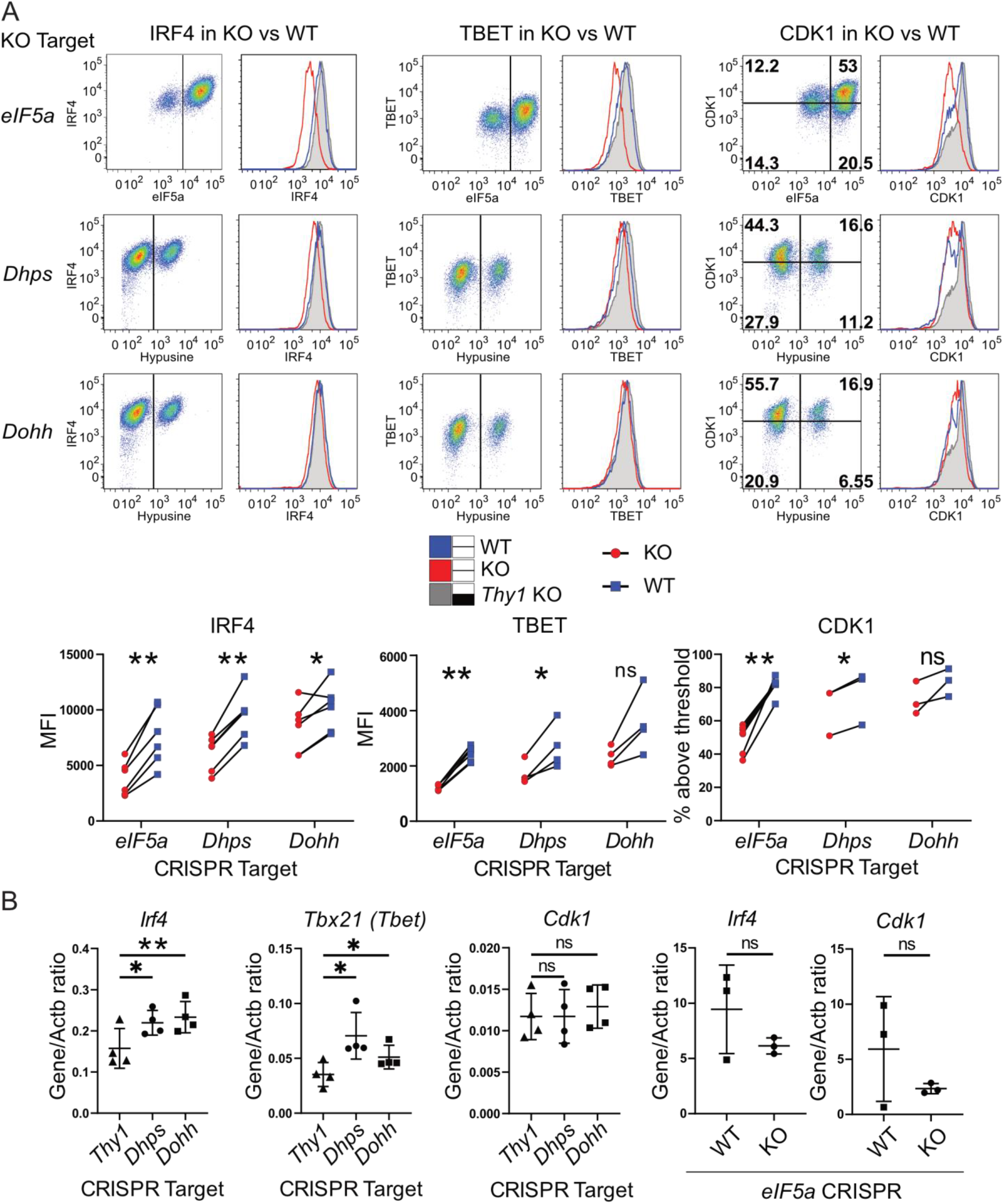
Flow cytometry validation of eIF5a regulated proteins. (A) Intracellular stains for eIF5a or hypusine were used to identify WT and KO *eiF5a*, *Dhps*, or *Dohh* CRISPR cells (representative dot plots on left). Overlay histograms on right show IRF4, TBET and CDK1 expression in electronically gated WT and KO samples with quantification of multiple paired samples shown below (n ≥ 3, *. P < 0.05, **. P < 0.01, paired T test). (B) mRNA of these genes were quantified by RT-qPCR in fresh bulk transfected cells (*Thy1*, *Dhps*, and *Dohh*) or fixed sorted cells for eIF5a KO (n ≥ 3, *. P < 0.05, **. P < 0.01, unpaired T test).

### Spermidine availability affects some but not all eIF5a-regulated proteins

The polyamine spermidine is produced naturally in cells and its abundance decreases with age. Exogenous supplementation of spermidine has been shown to increase organismal longevity (Eisenberg et al., 2009) and restore age-related T cell functions in an autophagy-dependent manner (Puleston et al., 2014). In addition to being required for hypusination of eIF5a, spermidine is also known for inhibiting the acetyltransferase activity of Ep300 and thereby inducing autophagy (Pietrocola et al., 2015). This non-eIF5a-dependent activity of spermidine may have contributed to discrepancies between cellular phenotypes of GC7 (a spermidine derivative) treated cells and *Eif5a/Dhps* KO cells. To further understand how spermidine affects T cell function via eIF5a, activated T cells were treated with an ornithine decarboxylase inhibitor DFMO for 48h to halt spermidine biosynthesis. Additionally DMFO-treated cells were supplemented with exogenous spermidine or were treated with GC7.

Quantitation of cellular hypusine levels reflected the predicted outcome of these treatments. The DFMO-treated cells had decreased hypusine, which was restored to a level comparable to untreated cells by spermidine supplementation. Depletion of hypusination by DFMO was exacerbated by additional GC7 treatment (Fig 6A). However, when we examined the abundance of total and selected proteins whose production we identified as being sensitive to loss of eIF5a by flow cytometry, an altered pattern emerged (Fig. 6B). Puromycin incorporation and CDK1, IRF4, TBET, and IFNγ abundance decreased following DFMO treatment, and were restored by exogenous spermidine addition, indicating they are very sensitive to cellular spermidine levels. Addition of GC7 further decreased Puromycin incorporation, CDK1 and IFNγ expression but did not impact significantly on the abundance of TBET and IRF4 protein. Interestingly, although IFNγ production was affected by DFMO, as reported recently for CD8 T cells (Alsaleh et al., 2020) and DFMO+GC7, the protein was still detectable in all cells, which was in contrast to *Eif5a/Dhps* KO cells where a significant proportion of the population produced no detectable IFNγ (Fig 2B). In contrast, TNFα, CD25 and CDC45 were only decreased in DFMO+GC7-treated cells and not following treatment with DFMO alone, indicating that they are less sensitive to spermidine depletion, despite showing some regulation by eIF5a (Fig 2B and Table S1).

**Fig 6.**
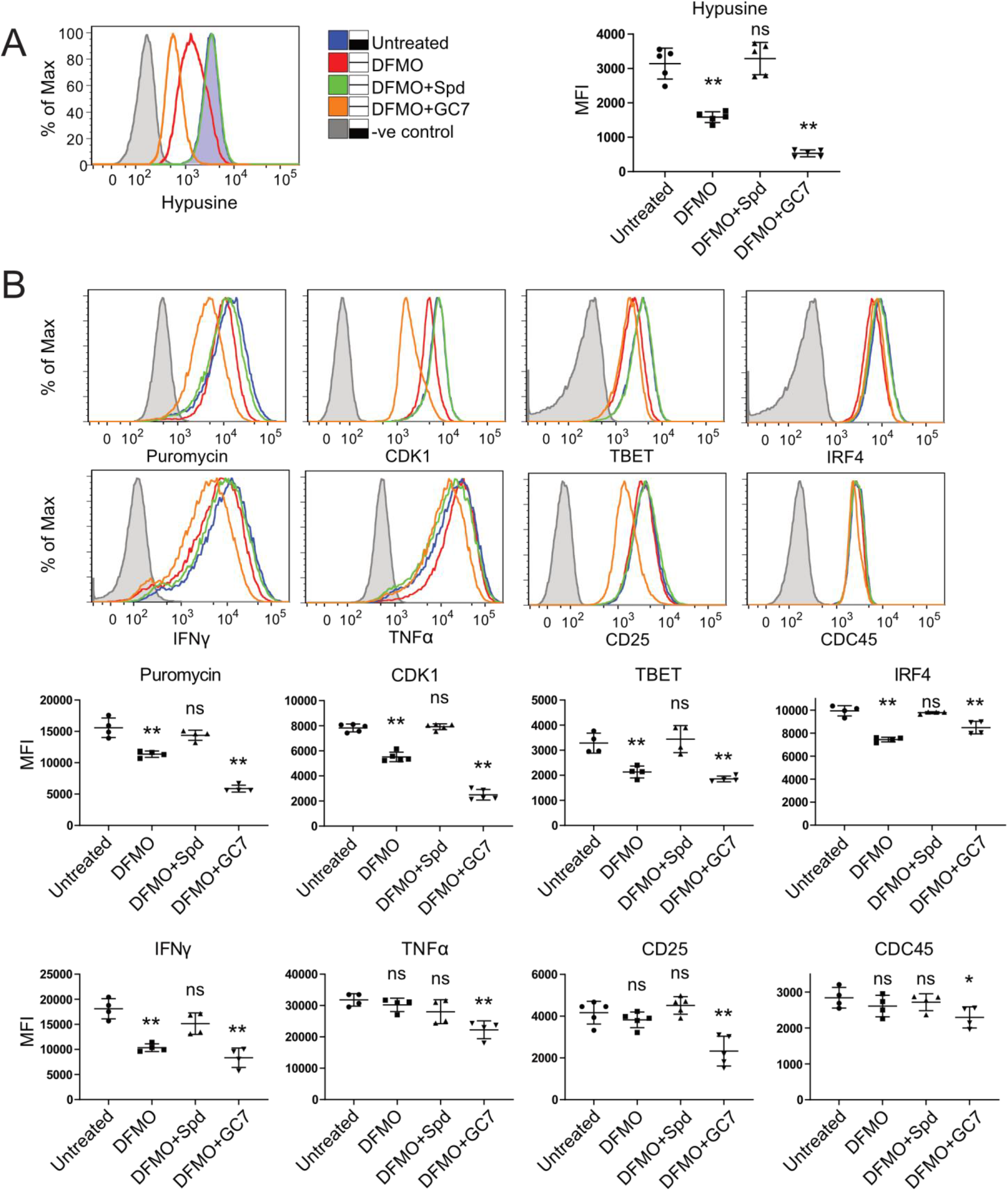
Restricting availability of polyamine spermidine to cells decreased eIF5a hypusination and some but not all eIF5a-regulated genes. (A-B) OT-1 cells were stimulated with peptide for 2d and cultured in IL-2 for a further 2d before treating with IL-2 only, 0.5mM DFMO, DFMO + 10μM spermidine, or DFMO + 10μM GC7 for another 2d. Cells were restimulated with peptide for a final 4h and labelled with puromycin for 10 mins and then harvested for intracellular staining and flow cytometry. (A) Inhibition of spermidine biosynthesis by DFMO reduces hypusine staining which is rescued by exogenous supplement of spermidine or further impaired by treatment with GC7. MFIs of hypusine from 5 independent experiments are plotted on the right (**p < 0.01, unpaired T test). (B) The same samples were stained for cytokines (IFNγ and TNFα), transcription factors (IRF4 and TBET), cell cycle regulators (CDK1 and CDC45), CD25 (IL2 receptor α), and puromycin. MFIs from ≥4 independent experiments were shown in the lower panel. Statistic tests were performed against cells treated with IL-2 only (*p < 0.05, **p < 0.01, unpaired T test).

It is likely that that GC7 treatment incompletely blocked eIF5a hypusination and, therefore, function and so had a milder effect on the down-regulation of IRF4 and TBET compared to *Eif5a* or *Dhps* KO (Fig S3). Together these data indicate that the maintenance of some cellular proteins such as CDK1, TBET and IRF4 are very sensitive to the availability of functional hypusinated eIF5a. On the other hand, other proteins such as the cytokine IFNγ, requires total ablation of eIF5a or DHPS in order to show a profound reduction in abundance. We could further demonstrate eIF5a-independent effect of GC7 on CD25 (IL-2ra), a crucial receptor for T cell expansion and survival, which was only mildly reduced in *Eif5a* KO or following DFMO treatment, but significantly down-regulated by DMFO+GC7 treatment (Fig 6B and S3).

## DISCUSSION

Our current understanding of eIF5a function largely comes from studies in yeast and transformed cell lines which proliferate continuously and therefore constitutively express active hypusinated eIF5a. In comparison, little has been reported on how eIF5a is regulated in primary mammalian cells. It was interesting therefore, that in primary T cells that transited between a naïve and an activated phenotype, a significant regulation of eIF5a activity occurred post-translationally. We demonstrated that naïve murine CD8 T cells have abundant expression of eIF5a protein, but proportionately less displayed hypusine modification compared to that found in activated T cells. Upon T cell activation, eIF5a becomes hypusinated in a dynamic fashion by the transcriptional upregulation of DHPS and DOHH, the enzymes dedicated to the transfer the 4-aminobutyl moiety from spermidine onto Lys^50^ of eIF5a. Similarly, human peripheral blood lymphocyte were shown to increase eIF5a hypusination upon activation with mitogens (Cooper et al., 1982). Using CRISPR knockout of the modifying enzymes or eIF5a itself, we confirmed that in T cells hypusination was important for eIF5a function. However the phenotype of the eIF5a knockout was more extreme than that of DHPS or DOHH knockouts suggesting that unmodified eIF5a might have some residual activity, although we are unable to exclude that a fraction of hypusinated eIF5a protein is very long-lived and that even small lingering quantities can still promote translation of some proteins. We also found that the deoxyhypusinated intermediate form of eIF5a had clearly detectable, albeit reduced functionality in T cells, as knockout of DOHH, which catalyses the last irreversible step in the conversion of lysine to hypusine, had much less impact than the removal of DHPS. Two key pathways in T cells which we could verify were primarily controlled at the level of translation by functional eIF5a, were control of cell cycle and cytokine production, both essential for T cell function.

Controlling protein production by regulating polypeptide elongation is emerging as an important means of translational control beyond that exerted by the more extensively studied regulators of translation initiation (Richter and Coller, 2015). Dysregulation of elongation has also been shown to contribute to diseases such as neurodegeneration and cancer (Knight et al., 2020; Richter and Coller, 2015). The availability of aminoacyl-transfer RNAs and the action of key proteins such as eEF1A, eEF2 and eEF2K have been well documented in controlling the rate of polypeptide formation (Choi et al., 2018). In contrast the role of eIF5a is less well understood but it is thought to be particularly important for elongation through certain motifs in the mRNAs particularly those encoding proline-proline and other charged residues (Schuller et al., 2017). We confirmed that eIF5a regulates de novo translation of subsets of cellular proteins in T cells which are enriched in PPP motifs, as previously described (Gutierrez et al., 2013), although more than 50% of our identified eIF5a-regulated genes did not have these motifs in their coding sequences. Strong ribosome pausing has been observed in eIF5a-depleted yeast at non-proline, charged amino acid containing motifs such as DVG, DDG, GGT, and RDK (Schuller et al., 2017), although of these motifs, only DVG seemed slightly, but not significantly, enriched in primary T cells. Online motif discovery tools such as MEME Suite (Bailey et al., 2009) were not able to identify enriched motifs in the CDS of eIF5a-regulated genes compared to all expressed proteins. The mechanism for down-regulation of non-polyproline containing eIF5a-regulated proteins remains unresolved. One possibility is that those mRNAs are disadvantaged by changes in ribosome composition (Ferretti et al., 2017; Shi et al., 2017) which were suggested by alterations in nascent polypeptide abundance of ribosome associated proteins in eIF5a KO cells and which might further influence prioritisation of mRNA translation. In this case some of the more global down-regulation of translation may not be linked to the elongation activity of eIF5a directly but rather indirectly through its influence on ribosomal protein translation. Interestingly in this context is a recent description of a ribosomopathy that presents as immune deficiency and impaired T cell activation (Robertson et al., 2021), providing further evidence that successful CD8 T cell activation may be particularly sensitive to ribosome composition.

An important consequence of T cell activation is their differentiation to effector cells whose function requires cytokine secretion in a temporally and spatially regulated manner. It has been appreciated for some time that cytokine production in immune cells is controlled both transcriptionally and translationally, with the latter involving both miRNAs (Asirvatham et al., 2009) and RNA binding proteins (Anderson, 2010; Salerno et al., 2020). We found the production of some effector cytokines, including IFNγ, to be particularly sensitive to the availability of functional eIF5a. Our data indicate that eIF5a provides an additional layer of cytokine regulation able to influence both translation of cytokines directly and indirectly through translation of their transcription factors such as TBET and IRF4. In this context there is an interesting correlation between the decline of cytokine production, T cell proliferation (Sievert et al., 2014) and memory T cell formation (Puleston et al., 2014) observed in ageing and an age-related decline in eIF5a protein abundance in lymphocytes (Zhang et al., 2019a). Not only is eIF5a protein reduced but spermidine availability also declines with age (Eisenberg et al., 2009) and dietary spermidine supplementation in old mice has been shown to promote and improve T memory cell formation (Puleston et al., 2014) and B cell antibody production (Zhang et al 2019). Although indirect, these publications support an active role for eIF5a in maintaining T cell functionality in ageing.

A recent study examined the role of polyamine metabolism on the differentiation of CD4 T cells and showed that either T cell-specific loss of ornithine decarboxylase, which is essential for spermidine production, or loss of DOHH strongly biased differentiation of naïve CD4 T cells to becoming excessive IFNγ producers at a loss of Th17 cytokine production (Puleston et al., 2021). This effect was manifest at the level of chromatin remodelling which is important during naïve CD4 T cell differentiation. Although these data may initially appear contradictory to some of the results reported here, it is important to stress that our study looked at the influence of eIF5a on CD8 T cells in short term cultures that had been activated and differentiated prior to Crispr treatment. Indeed loss of eIF5a function resulted in death of the knockout population over a matter of days in our system. Our purpose was to document eIF5a translationally regulated proteins in acutely activated T cells and avoid any confounders associated with lineage differentiation, while the Puleston et al study was concerned with the impact of these pathways on long term differentiation and immune function in vivo.

Inhibition of eIF5a hypusination with GC7 is in consideration as a treatment for several cancers due to its anti-proliferative activity (Coni et al., 2020) and ability to induce cell death (Schultz et al., 2018), so it was of interest to evaluate whether GC7 treatment mirrored the effects of genetic ablation of DHPS or eIF5a in T cells. GC7 produced a very different cellular phenotype from *Dhps* knockout, as has been shown for a fibrosarcoma cell line (Oliverio et al., 2014). As a structural derivative of polyamine spermidine, GC7 is known to compete with spermidine for binding to DHPS and has been widely used as an inhibitor for functional maturation of eIF5a. As spermidine has functions independent of eIF5a, such as inhibition of Ep300 activity, it is also likely that GC7 has comparable effects. CD25 has been previously shown to decrease following GC7 treatment in CD4 T cells (Colvin et al., 2013) as was seen here in CD8 T cells. However knockout of *eIF5a* did not reduce CD25 to the same extent, suggesting GC7 at least partially reduces CD25 expression independently of eIF5a. In our LC-MS dataset, GC7 treatment resulted in 3044 down-regulated nascent proteins, while eIF5a KO decreased 369 proteins. These observations indicate a substantial inhibitory effect of GC7 that does not act through eIF5a. Furthermore, proteins down-regulated exclusively after GC7 treatment were not enriched in polyproline motifs, and they clustered significantly to GO terms associated with ribosome constituents, suggesting GC7 reduces cellular translation in a more global manner than eIF5a. In contrast, eIF5a knockout resulted in increased rather than decreased abundance of several ribosomal proteins.

Our data support the importance of eIF5a in regulating immune cell function. Targeting eIF5a hypusination has been shown to have anti-inflammatory effects in macrophages (Oedem Paulo de et al., 2014) and to regulate macrophage mitochondrial respiration and activation (Puleston et al., 2019). Our current study also supports a role for eIF5a in influencing cytokine production and proliferation in T cells both through directly regulating translation of subsets of mRNAs and indirectly by influencing the production of key transcription factors. Together these studies lend weight to the existence of eIF5a directed RNA regulons important for modulating immune responses.

## Supporting information

Key resources table

## Acknowledgements

This research was funded by the Wellcome Trust Investigator Award WT205014/Z/16/Z to RZ and by the Wellcome Trust/Royal Society Sir Henry Dale Fellowship 20611/Z/17/Z to TL.

We would like to thank the following at the University of Edinburgh: Dr Martin Waterfall IIIR for cell sorting expertise, the BVS animal facility in Ashworth labs and CRM for support with animal husbandry, and the proteomics facility at the Wellcome Centre for Cell Biology for mass spec. We also thank Dr Alehandro Brenes Murillo at the University of Dundee for valuable discussion and help regarding dataset analysis.

## Author contributions

Conceptualization: TT & RZ; Methodology: TT, VK and TL; Investigation: TT, XZ and VK; Formal analysis: TT; Data curation: TT and VK; Writing – original draft: TT and RZ; Writing review and editing: TT, TL and RZ; Funding acquisition: TL and RZ; Resources: RZ; Supervision: RZ.

## Online data deposition

Original and processed RNASeq dataset have been deposited in the Gene Expression Omnibus (GEO) database, https://www.ncbi.nlm.nih.gov/geo (accession no. GSE168731).

Original files for the nascent proteomic dataset are available at EBI PRIDE database, https://www.ebi.ac.uk/pride/ (accession no. PXD021063).

## Methods

### LEAD CONTACT AND MATERIALS AVAILABILITY

Requests for resources and reagents should be directed to and will be fulfilled by the Lead Contact, Rose Zamoyska.

### EXPERIMENTAL MODEL

Mice used in experiments were of C57BL/6 background strain carrying homozygous OT-1 transgenic T cell receptor on a Rag-1KO background. All mice were bred at the animal facility in the University of Edinburgh in accordance with the UK Home Office and local ethically approved guidelines. Mice of both sexes were used.

### METHOD DETAILS

#### Cell culture and stimulation

Lymph node (LN) OT-I T cells were cultured in IMDM medium (Sigma) supplemented with 10% FCS, L-glutamine, 100U/ml penicillin, 100U/ml streptomycin and 50μM 2β-ME. 100nM SIINFEKL (N4) peptide (Cambridge Peptides) were added to culture media and the cells were incubated at 37°C for the duration specified in each experiment.

For GC7 treatment for transcriptomic and nascent proteomic profiling, 2-day cultured activated cells were washed and then cultured for 2 days in culture media supplemented with 20ng/mL recombinant human IL-2 (PeproTech) in the presence or absence of 10μM GC7 (Merck Chemicals). For DFMO experiment, activated cells expanded for 2 days in IL-2 were cultured for additional 2 days in fresh IL-2, or IL-2 supplemented with 0.5mM DFMO (Bio-Techne), DFMO+10μM spermidine (Sigma-Aldrich), or DFMO+10μM GC7.

#### CRISPR-Cas9 transfection

The following guide RNA sequences were purchased from Integrated DNA Technologies: *eIF5a*(TGCTCAGCATTACGTAAGAA and CTTCGAGACAGGAGATGCAG), *Dhps* (ATACCTCGTGCAGCACAACA), *Dohh* (GCAGTATTCTACGGACCCAG), *Thy1* (ACAGACAAGCTGGTCAAGTG), and TRAC guide RNA (CAGGGTTCTGGATATCTGT). Cas9 ribonucleoprotein complexes were formed by mixing 2μL of 100μM tracrRNA and 2μL of 100μM crRNA in 25μL Nuclease-free duplex buffer (Integrated DNA Technologies, USA), incubating at 95°C for 5 min and then mixed with 2μL of 5mg/mL Truecut Cas9 protein (Thermo Fisher Scientific, USA) for 10 min at 37°. The RNP complexes were made immediately before the transfection and mixed with 10^6^ 2-day activated OT-1 cells. Transfections were performed with the Neon transfection system (Thermo Fisher Scientific, USA) following manufacturer’s instructions, with the setting 1600V, 10ms, 3 pulses. Transfected cells were cultured in IL-2-containing media for 3 days for *eIF5a* KO, and 4 days for *Dhps* and *Dohh* KO.

#### AHA-labelling and cell sorting

Cells were transferred to IL-2-containing RPMI media in the absence of methionine, at a concentration of 5 × 10^6^ cells/mL. Cells were starved of methionine for one hour then 4-Azido-L-homoalanine HCl (L-AHA) (Jena Bioscience) was added to the media (final concentration 40μM) and incubated for 2 hours at 37°C. For *eIF5a* CRISPR experiment, cells were fixed in 2% formaldehyde for 20 min and permeabilised with ice-cold 90% methanol for 1 hour. Staining was done using anti-eIF5a-C-terminus antibody (Abcam) and anti-rabbit secondary antibody at AlexaFluor 647 (Thermo Fisher Scientific). Sorting was undertaken with a FACS Aria (BD).

#### Click chemistry and preparation for LC-MS

Proteins from freshly frozen cell samples (12.5 × 10^6^ of non-AHA-labelled, 32.5 × 10^6^ of GC7 and 12.5 × 10^6^ of Mock, 4 samples each) were extracted with CHAPS Lysis buffer from the Click Chemistry Capture Kit (Jena Bioscience). Input cell numbers were adjusted to give similar amount of total injected peptides, ensuring similar detection coverage. For sorted fixed cells, proteins from 10 × 10^6^ cells were extracted using 310μL of 200mM Tris, 0.5% SDS, 200mM NaCl, pH8.0, with brief sonications to disturb the cell pellet, followed by reverse crosslinking at 95°C for 45 min and cooled on ice, and then mixed with equal volume of 16M Urea giving a final concentration of 8M Urea. Extracted proteins were incubated overnight with Alkyne agarose beads (Jena Bioscience), and downstream preparations were done using Click Chemistry Capture Kit following the manufacturer’s guidelines, followed by desalting over C18 tips (Thermo Fisher Scientific). GC7 and Mock cells were pooled for the generation of a high pH reverse phase fractionated peptide library. Cells were lysed in PBS, 2% SDS, cOmplete protease inhibitors (Sigma), and 20 U of Benzonase (Millipore), reduced with 25mM TCEP, and alkylated with 25mM iodoacetamide. 500μg protein was precipitated with 80% acetone overnight at −20°C, washed twice with 100% acetone and once in 90% ethanol. Pellets were digested with 1:50 wt/wt trypsin in 100mM triethylammonium bicarbonate at 37°C overnight, then desalted over a C18 Sep-Pak cartridge (Waters). Peptides were separated over a 4.6mm XBridge BEH C18 column (Waters) in 10mM ammonium formate (pH 9) and eluted over a linear gradient from 2-50% acetonitrile at 1 ml/min. 72 fractions were collected and concatenated into 12 fractions and dried. Peptide samples were resuspended in 0.1% TFA, half of the click chemistry captured peptide and approximately 0.5 μg of the 12 library fractions were injected for LCMS analysis. An Ultimate 3000 RSLCnano HPLC (Dionex, Thermo Fisher Scientific) was coupled via electrospray ionisation to an Orbitrap Elite Hybrid Ion Trap-Orbitrap (Thermo Fisher Scientific). Peptides were loaded directly onto a 75 μm x 50 cm PepMap-C18 EASY-Spray LC Column (Thermo Fisher Scientific) and eluted at 250 nl/min using 0.1% formic acid (Solvent A) and 80% acetonitrile/0.1% formic acid (Solvent B). Samples were eluted over 120 min stepped linear gradient from 1% to 30% B over 95 min, then to 45% B over a further 25 min. MS1 scans were acquired at 120k resolution over 350-1700 m/z and a ‘lock mass’ of 445.120025 m/z was used. This was followed by 20 data-dependent MS2 CID events in the ion trap at rapid resolution with a 2 Da isolation width, 5E3 target ion accumulation, 50 ms maximum fill time, a normalised collision energy of 35, a requirement of a 10k precursor intensity, and a charge of 2+ or more. Precursors within 5 ppm were dynamically excluded for 40 sec.

#### Systematic Dataset handling

Pair-end RNA sequencing dataset reads were generated by Novogene (GC7- and Mock-treated samples, 4 biological replicates) and were trimmed of low-quality and adapter sequences. Reads were aligned to the mouse genome (GRCm38.96) using STAR (Dobin et al., 2013) and mapped reads were counted using featureCounts (Liao et al., 2013). Differential expression analysis was performed with edgeR (Robinson et al., 2010).

Nascent proteomic data was analysed using MaxQuant version 1.6.2.6 (Cox and Mann, 2008). LC-MS/MS data was searched against the mouse reference proteome from UniProt derived from mouse genome GRCm38 (accessed on 5^th^ February, 2018), which contains 52,599 entries, allowing for 2 tryptic missed cleavages, allowing for variable methionine oxidation, protein N-terminal acetylation, and cysteine carbamidomethylation. The parameter “Individual peptide mass tolerance” was selected for variable precursor mass tolerances, with 0.5 Da or 20 ppm mass tolerances set for ion trap or orbitrap fragment ions, respectively. A target-decoy threshold of 1% was set for both PSM and protein false discovery rate. Match-between-runs was enabled with identification transfer within 0.7 min and a retention time alignment within 20 min window. Matching was permitted from the library parameter group, and ‘from and to’ the unfractionated parameter group, and second peptide search was enabled. iBAQ values were used for analysis. all values were normalised using mean of total iBAQ within sample groups to adjust for unequal sample loading. Since different number of input cells were used between groups to achieve similar coverage, the iBAQs were then normalised by the input cell numbers, followed by imputation where zero values were replaced by 1000 (Detailed in Fig S1A). Statistical tests were performed using Log2-transformed normalised iBAQ values. Unpaired T test with Benjamini-Hochberg correction (Benjamini and Hochberg, 1995) was used to determine significantly down-regulated proteins between GC7- and Mock-treated samples. Paired T test was used for eIF5a KO versus WT samples.

#### Flow cytometry

Cells were labelled with LIVE/DEAD™ Aqua Dead Cell Stain Kit (Invitrogen) and then fixed in Intracellular Staining Fixation Buffer (BioLegend) and permeabilised in Intracellular Staining Permeabilization Wash Buffer (BioLegend). Antibodies against eIF5a (EP527Y, Abcam), hypusine (Hpu24, Creative Biolabs), IFNg (XMG1.2, BioLegend), TNFa (MP6-XT22, eBioscience), IRF4 (3E4, eBioscience), TBET (4B10, BioLegend), CDK1 (A17, Abcam), CD25 (PC61, BioLegend), CDC45 (EPR5759, Abcam) and puromycin (12D10, Millipore) were used; for cell cycle analysis, fixed-permeabilised cells were stained with Hoechst 33342 (Sigma) and Ki-67 antibody (BD Biosciences). Cells were incubated with primary antibodies for 16 hours at 4°C, washed, followed by staining with Goat anti-rabbit Alexa Fluor 647 secondary antibody (Thermo Fisher Scientific) for 1 hour at RT. All analyses were done on singlet live events.

All acquisitions were done on a MACSQuant flow cytometer (Miltenyi Biotec).

#### Protein extraction and Western blotting

Cells were lysed in RIPA Buffer (Thermo Fisher Scientific) supplemented with a protease inhibitors cocktail (Sigma). The lysates were centrifuged to remove the debris. Reducing Laemmli buffer was added to the lysates, and samples were heated to 95°C and separated by SDS-PAGE. Proteins were transferred to Immobilon-FL PVDF membrane (Millipore), and membranes were incubated in Odyssey blocking buffer (LI-COR Biosciences) prior to incubation with the primary antibodies. Quantitative signals were generated with secondary Abs: IRDye® 680RD Goat anti-Rabbit and IRDye® 800CW Goat anti-Mouse (LI-COR Biosciences), and visualised using the Odyssey Infrared Imaging System (LI-COR Biosciences).

The following primary Abs were used: anti-ZAP70 (BD Biosciences), DHPS (ab224134, Abcam), eIF5a (EP527Y, Abcam), and hypusine (Hpu24, Creative Biolabs).

#### Transcript analyses by RT-qPCR

Total cellular RNA was extracted using Direct-zol RNA Miniprep kit (Zymo Research) and reverse transcribed using Lunascript kit (New England Biolabs). qPCR was performed on a Light Cycler 480 (Roche) using Brilliant III SYBR® Master Mixes (Agilent). Transcription level of each gene was normalised against the quantity of beta actin (Actb) in the sample. For fixed sorted cells, total cellular RNA was isolated using RecoverAll™ Total Nucleic Acid Isolation Kit (Thermo Fisher Scientific) and analysed as above.

#### QUANTIFICATION AND STATISTICAL ANALYSIS

Prism software (GraphPad) was used for statistical analyses. All data points represent the measurement of biological replicates. Unpaired or paired two-tailed Student’s t test was used for comparisons between two normally distributed datasets with equal variances unless specified. One-tailed Fisher’s exact test was used for testing enrichment of amino acid trimers. In the GC7-Mock nascent proteome dataset, Benjamini–Hochberg method was used to adjust P values of a family of multiple t tests with a single null hypothesis. P value was used to quantify the statistical significance of the null hypothesis testing. *p ≤ 0.05, **p ≤ 0.01.

## DATA AND CODE AVAILABILITY

Original data and code are available at https://doi.org/10.17632/vhyhnn96hf.1

## Supplementary Tables as Excel files

**Table S1: Normalised iBAQ values of nascent proteomes, differential expression analysis, and selection of proteins for presentation**

Legend: iBAQ values from LC-MS for GC7-treated, Mock-treated, eIF5a-KO and eIF5a-WT cells were normalised using average sum within sample groups, and then by the input cell numbers. Statistical tests were performed using Log2-transformed normalised iBAQ values. Unpaired T test with Benjamini-Hochberg correction was used to determine significantly down-regulated proteins between GC7- and Mock-treated samples. Paired T test was used for eIF5a KO versus WT samples.

**Table S2: Differential expression analysis of the RNASeq dataset between GC7- and Mock-treated cells, with top ten enriched KEGG pathways in significantly up- and down-regulated genes.**

Legend: The dataset was trimmed of low-quality and adapter sequences. Reads were aligned to the mouse genome (GRCm38.96) using STAR and mapped reads were counted using featureCounts. Differential expression analysis was performed with edgeR. Pathway enrichment analysis was conducted with WebGestalt.

**Table S3: Determination of genes translationally regulated by eIF5a using Venn analyses.**

Legend: Genes translationally down-regulated upon GC7-treatment was defined as those not down-regulated in RNA levels but decreased in nascent peptide levels following the treatment. This list of genes was overlaid with nascent proteins down-regulated in eIF5a-KO cells and the common genes between them were defined as genes translationally regulated by eIF5a.

---

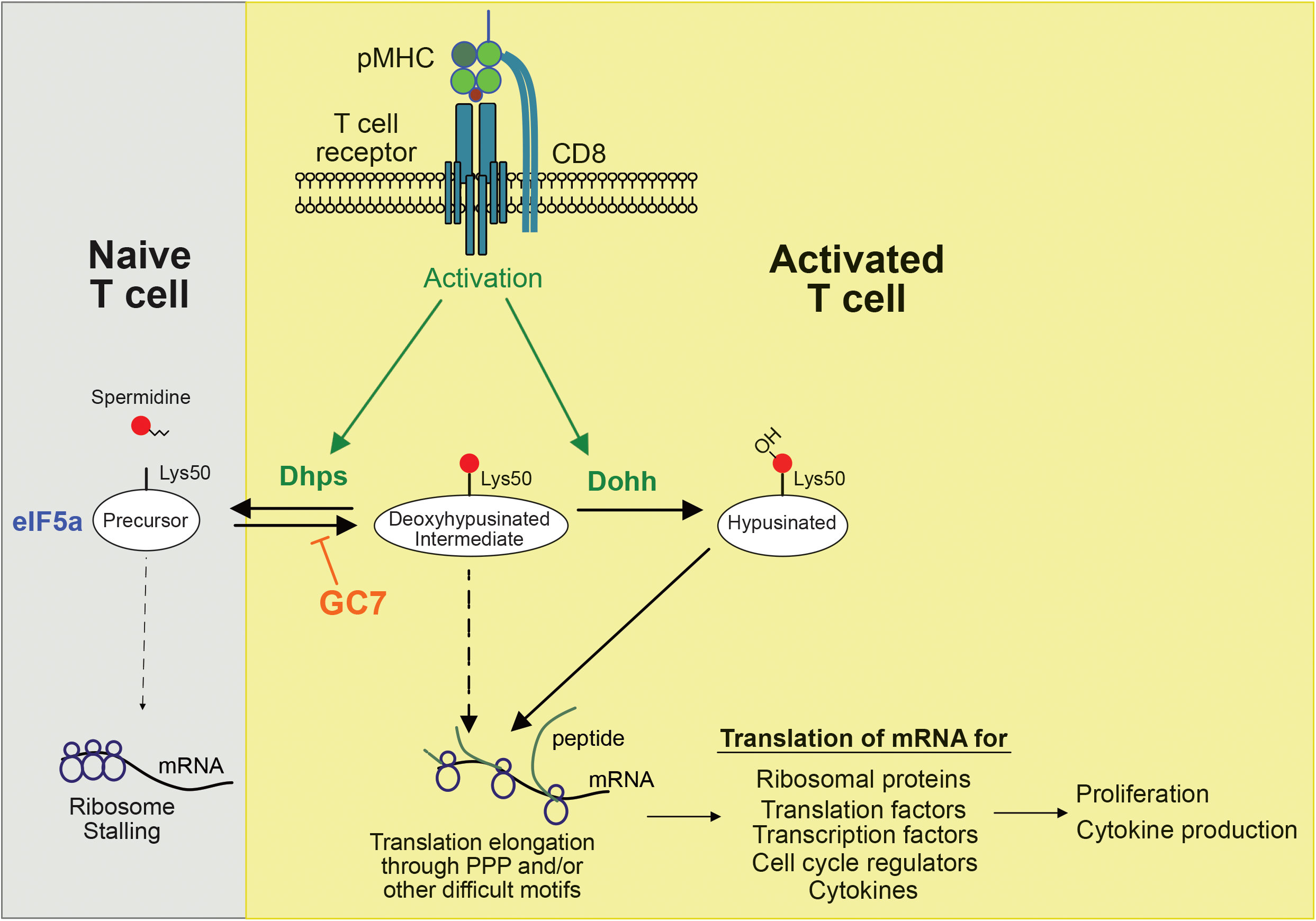
Graphical summary

**Fig S1.**
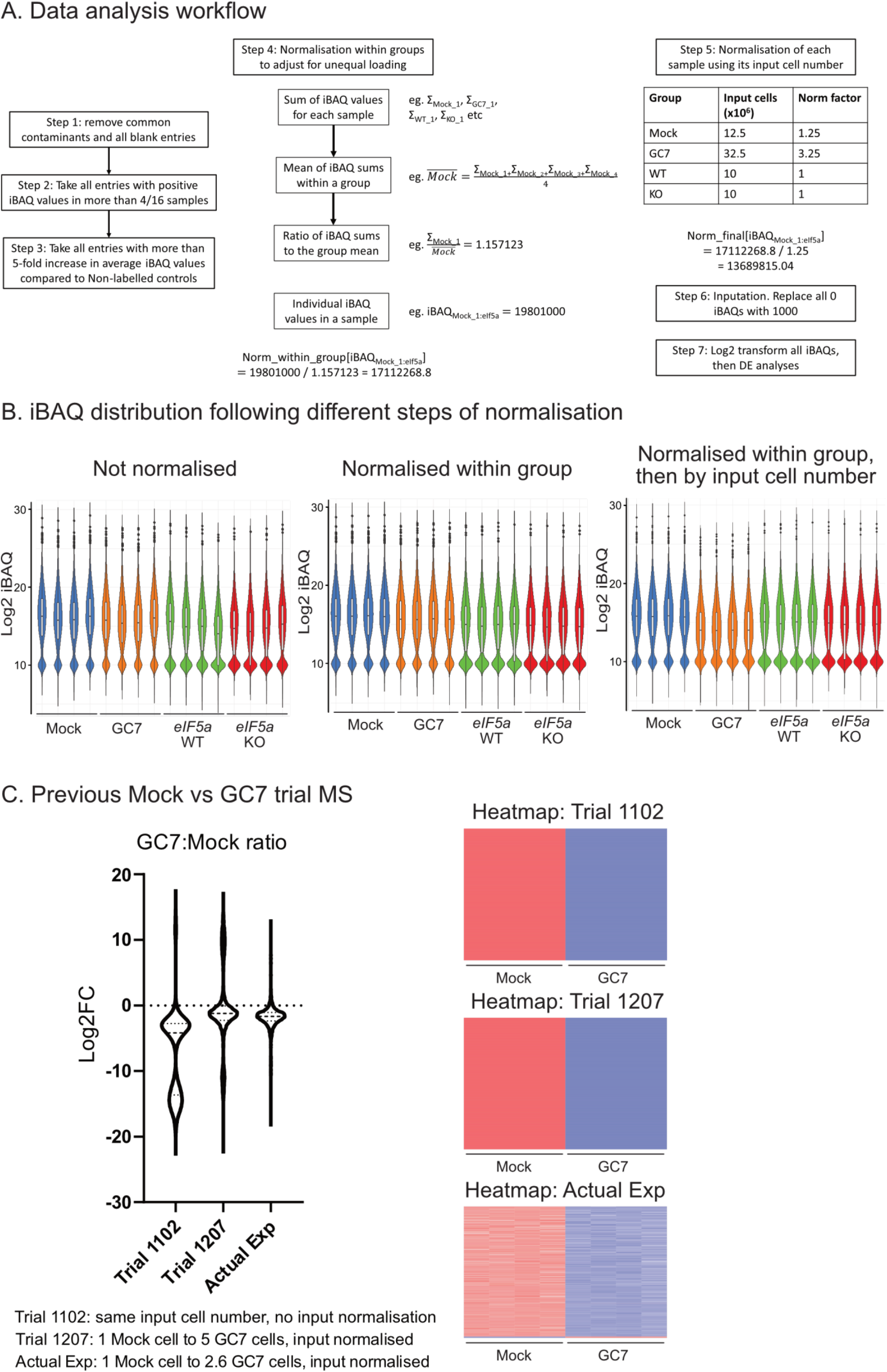
Normalisation of nascent proteome dataset, and comparison with previous trial experiments. (A) Workflow illustrating how iBAQ values were processed and normalised. (B) Violin plots showing Log2 values of iBAQ without any normalisation, after normalisation within group, and then after normalisation by input cell number. (C) Violin plot showing Log2 GC7:Mock ratios in two previous trial experiments (1102: same number of input cells between Mock- and GC7-treated samples; 1207: 1 Mock cell to 5 GC7 cells) and in the actual experiment reported here. Heat maps (Babicki et al., 2016) showing Log2 iBAQ of 2500 most different proteins in the three experiments are shown below.

**Fig S2.**
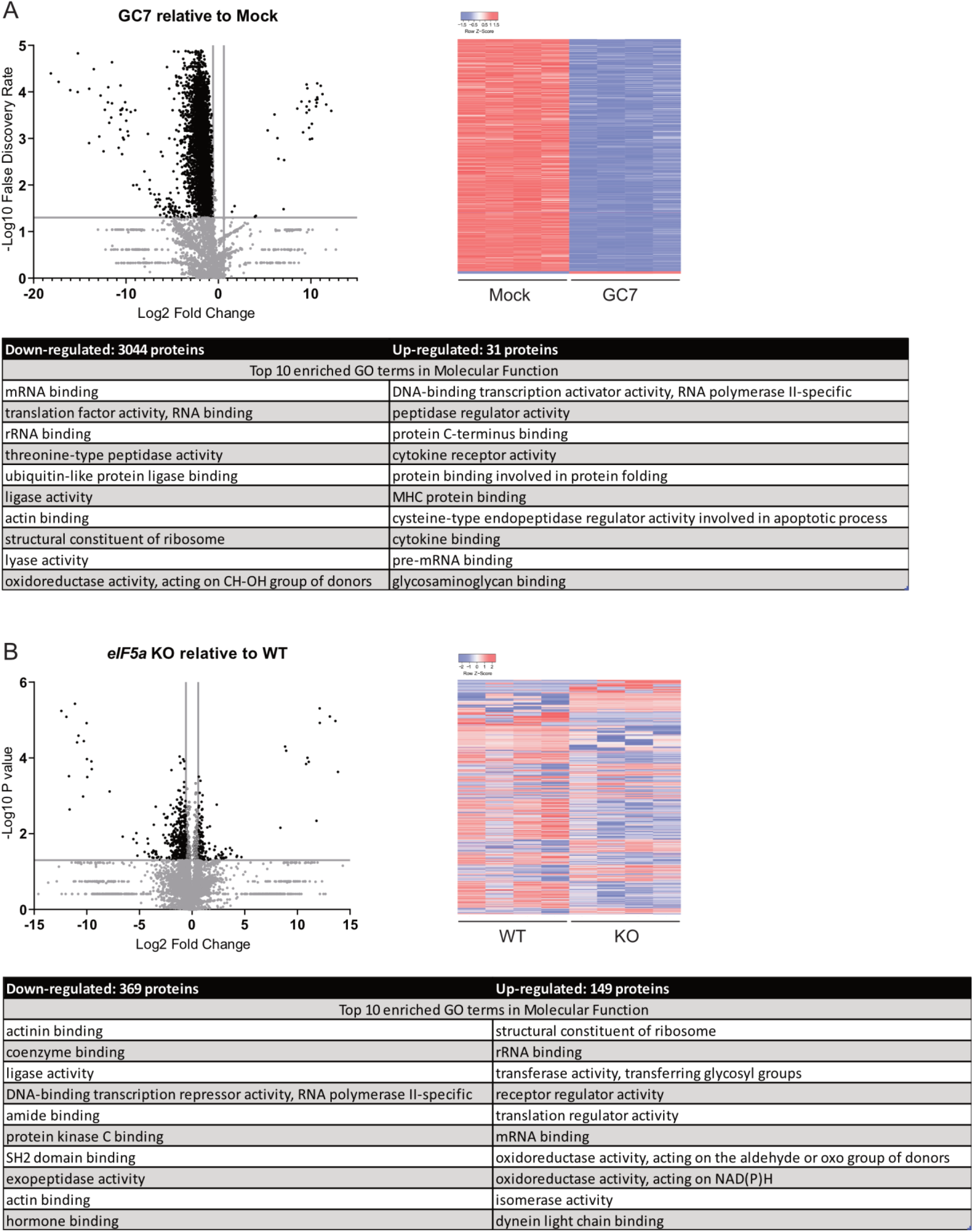
Analyses of nascent proteins up- or down-regulated following (A) GC7-treatment or (B) eIF5a KO. Volcano plots were plotted using the significance criterion described in methods. Heat maps were constructed using the top 2500 significantly changed proteins (Babicki et al., 2016), relative protein abundance is graded from low (blue), medium (white) to high (red) to allow comparisons between samples. Top 10 enriched GO terms in molecular function were generated using WebGestalt (Liao et al., 2019).

**Fig S3.**
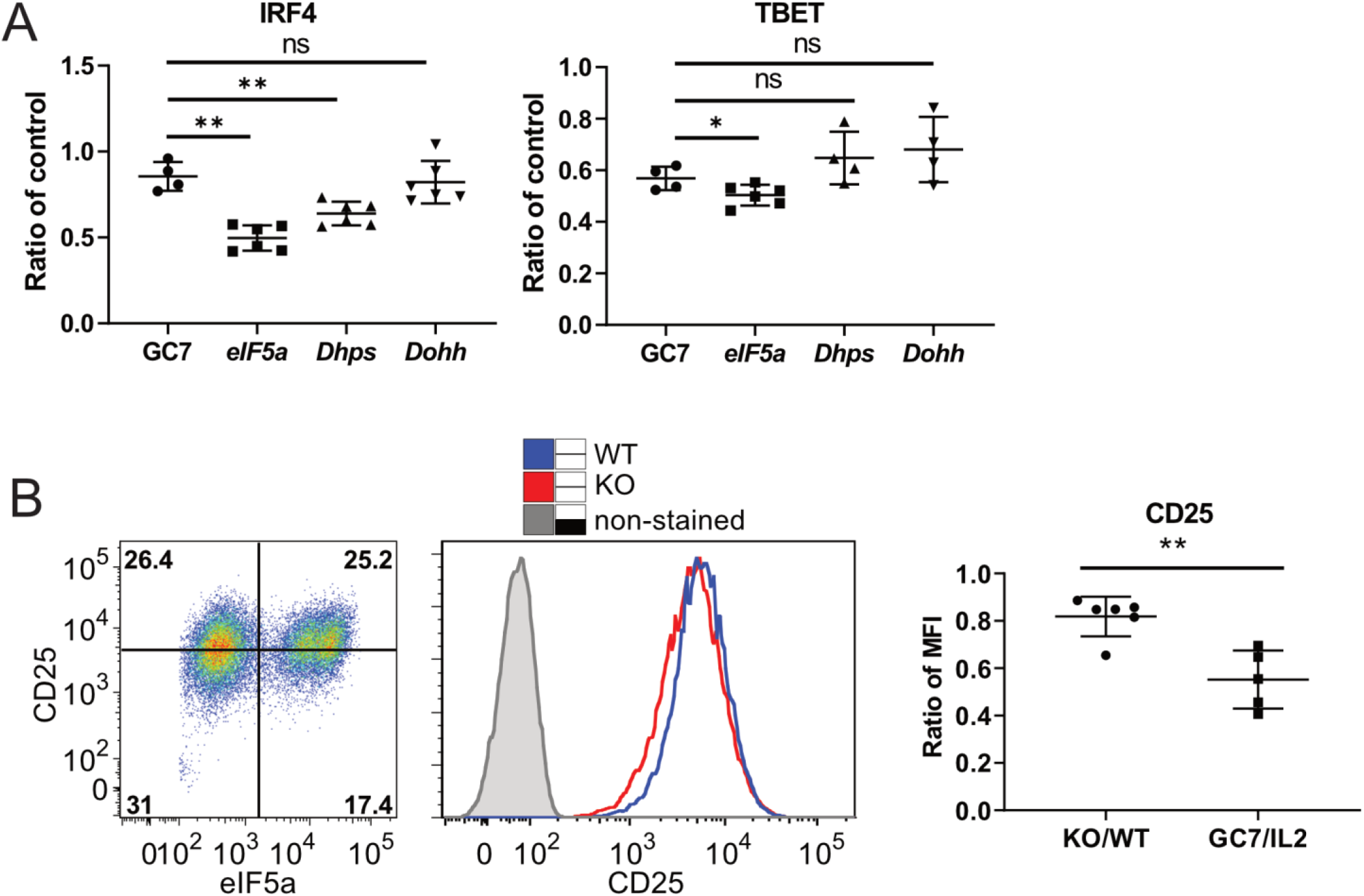
eIF5a KO and GC7 treatment have different impacts on expression of individual proteins. (A) MFIs for IRF4 and TBET staining were taken from Fig 3 and 4. To compare across experiments the ratio of treated to control (GC7 vs untreated or KO versus WT cells) were calculated and plotted (n≥4, *p < 0.05, **p < 0.01, unpaired T test). (B) An example of CD25 expression versus eIF5a measured by flow cytometry at Day 3 after CRIPSR-Cas9 transfection is shown in the dot plots and histogram overlays. The fold reduction of MFIs in KO from 6 independent experiments were compared to those of GC7 treated cells in Fig 4 (**p < 0.01, unpaired T test).

