## Supplementary material for "Essential role of translation factor eIF5a in cytokine production and cell cycle regulation in primary CD8 T lymphocytes": Key resources table

| REAGENT or RESOURCE | SOURCE | IDENTIFIER |
| --- | --- | --- |
| Antibodies | | |
| eIF5a clone EP527Y | Abcam | ab32407; RRID:AB_732132 |
| Hypusine clone Hpu24 | Creative Biolabs | PABL-202 |
| IFNg clone XMG1.2 (PE conjugated) | BioLegend | 505808; RRID:AB_315402 |
| TNFa clone MP6-XT22 (PE-Cy7 conjugated) | eBioscience | 25-7321-82; RRID:AB_11042728 |
| IRF4 clone 3E4 (eFlour 450 conjugated) | eBioscience | 48-9858-82; RRID:AB_2574135 |
| TBET clone 4B10 (Brilliant Violet 421 conjugated) | BioLegend | 644832; RRID:AB_2686976 |
| CDK1 clone A17 | Abcam | ab18; RRID:AB_2074906 |
| CD25 clone PC61 (PE-Cy7 conjugated) | BioLegend | 102016; RRID:AB_312865 |
| CDC45 clone EPR5759 | Abcam | ab126762; RRID:AB_11140216 |
| Puromycin clone 12D10 (Alexa Fluor® 488 conjugated) | Millipore | MABE343-AF488; RRID:AB_2736875 |
| Ki-67 clone B56 | BD Biosciences | 556027; RRID:AB_2266296 |
| Goat anti-rabbit IgG Alexa Fluor 647 | Thermo Fisher Scientific | A27040; RRID:AB_2536101 |
| ZAP70 clone 29 | BD Biosciences | 610239; RRID:AB_397634 |
| DHPS | Abcam | ab224134 |
| Goat anti-Rabbit IgG (IRDye® 680RD conjugated) | LI-COR Biosciences | 926-68071; RRID:AB_10956166 |
| Goat anti-Mouse IgG (IRDye® 800CW conjugated) | LI-COR Biosciences | 926-32210; RRID:AB_621842 |
| Chemicals, peptides, and recombinant proteins | | |
| LIVE/DEAD™ Aqua Dead Cell Stain Kit | Invitrogen | L34957 |
| Intracellular Staining Fixation Buffer | BioLegend | 420801 |
| Intracellular Staining Permeabilization Wash Buffer | BioLegend | 421002 |
| SIINFEKL (N4) peptide | Cambridge Peptides |  |
| Recombinant human IL-2 | PeproTech | AF-200-02-1mg |
| N1-Guanyl-1,7-diaminoheptane (GC7) | Merck Millipore | 259545-10MG |
| Difluoromethylornithine (DFMO) | Bio-Techne | 2761/50 |
| Spermidine | Sigma-Aldrich | S0266-1G |
| Nuclease-free duplex buffer for CRISPR guide RNA | Integrated DNA Technologies | 11-05-01-12 |
| Truecut Cas9 protein v2 | Thermo Fisher Scientific | A36499 |
| 4-Azido-L-homoalanine HCl (L-AHA) | Jena Bioscience | CLK-AA005-10 |
| Alkyne agarose beads | Jena Bioscience | CLK-1032-2 |
| RIPA Buffer | Thermo Fisher Scientific | 89900 |
| Protease inhibitors cocktail | Sigma-Aldrich | P8340 |
| Odyssey blocking buffer | LI-COR Biosciences | 927-40000 |
| bisBenzimide H 33342 trihydrochloride (Hoechst 33342) | Sigma-Aldrich | B2261 |
| Critical commercial assays | | |
| Click Chemistry Capture Kit | Jena Bioscience | CLK-1065 |
| Direct-zol RNA Miniprep kit | Zymo Research | R2051 |
| RecoverAll™ Total Nucleic Acid Isolation Kit for FFPE | Thermo Fisher Scientific | AM1975 |
| Lunascript RT SuperMix Kit | New England Biolabs | E3010L |
| qPCR Brilliant III SYBR Master Mix | Agilent | 600883 |
| Deposited data | | |
| RNASeq | Gene Expression Omnibus | GSE168731 |
| Nascent proteomes | EBI PRIDE | PXD021063 |
| Original data for figures and tables | Mendeley data | https://doi.org/10.17632/vhyhnn96hf.1 |
| Experimental models: Organisms/strains | | |
| Mouse: C57BL/6, Rag-1KO, OT-1 homozygous |  |  |
| Oligonucleotides | | |
| crRNA: eIF5a (TGCTCAGCATTACGTAAGAA and CTTCGAGACAGGAGATGCAG) | Integrated DNA Technologies | Custom synthesised RNA |
| crRNA: Dhps (ATACCTCGTGCAGCACAACA) | Integrated DNA Technologies | Custom synthesised RNA |
| crRNA: Dohh (GCAGTATTCTACGGACCCAG) | Integrated DNA Technologies | Custom synthesised RNA |
| crRNA: Thy1 (ACAGACAAGCTGGTCAAGTG) | Integrated DNA Technologies | Custom synthesised RNA |
| tracrRNA (CAGGGTTCTGGATATCTGT) | Integrated DNA Technologies | Custom synthesised RNA |
| RT-qPCR Primer: Actb (forward: ATGGAGGGGAATACAGCCC; reverse: TTCTTTGCAGCTCCTTCGTT) | Integrated DNA Technologies | Custom synthesised DNA |
| RT-qPCR Primer: IFNg (forward: GAGCTCATTGAATGCTTGGC; reverse: GCGTCATTGAATCACACCTG) | Integrated DNA Technologies | Custom synthesised DNA |
| RT-qPCR Primer: TNFa (forward: CTGAACTTCGGGGTGATCGG; reverse: GGCTTGTCACTCGAATTTTGAGA) | Integrated DNA Technologies | Custom synthesised DNA |
| RT-qPCR Primer: Irf4 (forward: TCCGACAGTGGTTGATCGAC; reverse: CCTCACGATTGTAGTCCTGCTT) | Integrated DNA Technologies | Custom synthesised DNA |
| RT-qPCR Primer: Tbx21 (Tbet) (forward: AACACACACGTCTTTACTTTCCA; reverse: CGTATCAACAGATGCGTACATGG) | Integrated DNA Technologies | Custom synthesised DNA |
| RT-qPCR Primer: Cdk1 (forward: AGAAGGTACTTACGGTGTGGT; reverse: GAGAGATTTCCCGAATTGCAGT) | Integrated DNA Technologies | Custom synthesised DNA |
| Software and algorithms | | |
| FlowJo | FlowJo (commercial) | https://www.flowjo.com |
| Prism | GraphPad (commercial) | https://www.graphpad.com/scientificsoftware/  prism/ |
| Image Studio Lite | LI-COR (commercial) | https://www.licor.com/bio/image-studio-lite/ |
| R studio | R Studio (commercial) | https://www.rstudio.com/ |
| MaxQuant | Max-Planck-Institute of Biochemistry | https://www.maxquant.org/ |
| Other | | |
| Neon transfection system | Thermo Fisher Scientific | MPK5000 |
| Orbitrap Elite™ Hybrid Ion Trap-Orbitrap Mass Spectrometer | Thermo Fisher Scientific |  |
